# Low-disturbance Farming Regenerates Healthy Deep Soil towards Sustainable Agriculture

**DOI:** 10.1101/828673

**Authors:** Fangbo Deng, Hongjun Wang, Hongtu Xie, Xuelian Bao, Hongbo He, Xudong Zhang, Chao Liang

**Author notes:** These authors contributed equally to this work (Fangbo Deng, Hongjun Wang, Hongtu Xie).

## Abstract

Intensive conventional farming has degraded farmland topsoil and seriously threaten food and environment security globally. Although low-disturbance practices have been widely adapted to restore soil health, whether this measure in a long run can potentially recover the critical deep soil to meet sustainable intensification of crop production are still unclear. Here we compared soil microbiome, physicochemical parameters along 3-m deep soil profiles, and crop yield in Northeast China subjected to ten years of farming practices at 3 levels of disturbance, including conventional tillage (CT), no-tillage without stover mulching (NTNS), and no-tillage with stover mulching (NTSM). We found that low-disturbance practices (NTNS and NTSM) promoted the ability of the deep soil to retain water, nitrogen and salt-extractable organic, regenerated whole-soil microbial diversity and metabolic function, improved topsoil organic carbon stock and corn yield in the drought year, showed the potential to reduce energy consumption and greenhouse gas emissions, thus regenerating highly efficient, sustainable agriculture.

## Introduction

Since the Industrial Revolution, the rate of soil carbon loss has increased dramatically, resulting in a global carbon debt due to agriculture of 116 Pg carbon for the top 2 m of soil(Sanderman *et al*., 2017). The loss of soil carbon in farmlands has not only changed global climate but also produced catastrophic cascade impacts on global food and environment security, as soil carbon is the cornerstone for healthy and productive soil that will be needed to sustainably feed 10 billion people in 2050 (United Nations, World Population Prospects 2019). It is well known that intensive conventional farms with high energy inputs (chemical fertilizers) and disturbance (e.g. tillage/compaction, burn/remove stover) have caused a series of environmental issues, like water pollution, biodiversity loss, freshwater depletion and climate change(Congreves *et al*., 2015). Even worse, increasing the amount of chemical fertilizer is unlikely to continue the increase in quantity and quality of food products worldwide(Vitousek *et al*., 2009), while the options to expand the farmland area at the expense of nature and biodiversity that already under pressure is limited(Springmann *et al*., 2018). Moreover, the topsoil disturbance, tillage in particular, prevents root growth into deeper soil(Kemper *et al*., 2011), thus critically minimizing the exploitation of deep soil profile(Billings *et al*., 2018) and reducing crop’s nutrient using efficiency and its resilience to drought.

Since the 1970s, low-disturbance practices (e.g. reduced tillage, no-tillage and stover mulching) have been gradually applied to restore soil health and reduce non-point source pollution(Salem). Growing evidence shows that no-tillage and stover mulching boosted top-soil organic carbon (SOC)(Blanco-Canqui and Lal, 2008; Liu *et al*., 2014), increased soil aggregate stability(Song *et al*., 2019) and reduced soil erosion and surface runoff(Prosdocimi *et al*., 2016). All these benefits from low-disturbance tillage practices are tied with complex microbial processes that contribute to soil structure formation, modulate the biogeochemistry cycling of nutrients and drive soil carbon transformation and stabilization(Schimel and Schaeffer, 2012; Bardgett and Van Der Putten, 2014; Liang *et al*., 2017). Microorganisms also respond quickly to changes in soil environment conditions, due to their high surface to volume ratio, which can provide an early signal in soil improvement or degradation, thus has been recommend as a key indicator of soil health(Nielsen *et al*., 2002). However, the effect of tillage practices on soil microbial communities are complex and diverse(Helgason *et al*., 2010; Sun *et al*., 2018; Li *et al*., 2020; Rincon-Florez *et al*., 2020) and most studies by now mainly focused on farmland topsoil or soils within 1-m depth(Hartman *et al*., 2018; Nevins *et al*., 2018; Alahmad *et al*., 2019), and soil below 1 meter, which contains more carbon than the topsoil(Jobbágy and Jackson, 2000) and belongs to Earth’s Critical zone, was often overlooked. However, microbes inhabiting in the deep soils (> 1 m) may substantially impact long-term carbon sequestration, mineral weathering and crop production(Eilers *et al*., 2012; Sagova-Mareckova *et al*., 2016; Pries *et al*., 2017), and play important roles in bridging aboveground vegetation with parent soils and even acts as an essential buffer protecting underground water(Chorover *et al*., 2007). Moreover, studies indicate many crops’ roots (depending on the species and management) can penetrate over one meter in depth(Stone *et al*., 2001; Thorup-Kristensen, 2006), which means they can potentially forage for nutrient and water in deep soil and impact deep-soil microbial community(Pierret *et al*., 2016; Thorup-Kristensen *et al*., 2020). More importantly, tillage practices significantly influence root growth(Qin *et al*., 2004; Kemper *et al*., 2011; Nunes *et al*., 2019), the change of surface soil carbon by land use or agricultural practice influenced the carbon dynamic in deep soil layers has also been observed(Fontaine *et al*., 2007; Tautges *et al*., 2019; Dal Ferro *et al*., 2020). We thus hypothesized that: 1) different tillage practices will affect the vertical distribution of the microbial community by altering the spatial distribution of nutrients and dissolved organic carbon, thereby changing the ecosystem function; 2) lowest disturbance practice-no-tillage with 100% stover mulching as a nature-based solution, close to undisturbed natural ecosystem, would regenerate healthy deep-soil with highly diversified and functional microbes over time toward a highly resilient, sustainable agricultural ecosystem.

Recent research shows that corn belts in the U.S.A., western Europe, and China have experienced the most soil carbon loss globally(Sanderman *et al*., 2017). The corn belt in Northeast China is considered as the “breadbasket” of the country, having the largest grain production and overlapping with the most fertile Mollisol region that sustains 3% of population in the world(Liu *et al*., 2010), accounting for over 30% of corn production of China(Liu *et al*., 2012). Here, a 10-year manipulative experiment was conducted at a temperate corn farm in Northeast China, investigating farming practices with three levels of disturbance: high disturbance—conventional tillage (CT), low disturbance—no-tillage without stover mulching (NTNS) and lowest disturbance—no-tillage with 100% stover mulching (NTSM) (described in the Methods and Fig. S1). We compared soil physicochemical properties, fine-root associated organic carbon, and microbial communities of the 3-m soil profiles at the end of dormant season (legacy effects of nutrients) after the 10-year manipulation, and multi-year corn yield as well. We aimed at testing our hypotheses and providing a general picture of soil microbial community in the entire soil profiles, which are important for comprehensive recognize the impact of low-disturbing tillage practices on ecosystem function and developing effective strategies to ensure the sustainable development of agriculture.

## Results

### Soil properties and corn yield

The SOC, TN and C/N ratio substantially decreased from the soil surface to around 150 cm depths and then remained unchanged within 150-300 cm (Fig. 1). The NTSM slightly increased SOC, TN and C/N ratio at 0-20 cm soil layers compared with the NTNS and the CT (Fig. 1 and Table S1). The NTSM and NTNS reduced soil pH in surface and deeper layers (Fig. 1d) and increased soil moisture at surface layers (0-60 cm) (Fig. 1e). In the CT plots, soil NO_3_^−^-N concentration first decreased and then increased remarkably, ranged from 4.19 to 23.32 mg kg^−1^ (Fig. 1i). However, under the NTNS and NTSM treatments, soil NO_3_ ^−^-N decreased significantly at 0-40 cm then increased to the maximum at 120-150 cm. Interestingly, above 120-150 cm layer, NO_3_^−^-N was significantly higher with low-disturbance practices than conventional tillage, while the soil below 150 cm under low-disturbance practices had much lower NO_3_^−^-N compared to conventional tillage (Fig. 1i). The NTNS plots contained much higher amounts of ammonium than the CT and the NTSM plots (Fig. 1h). Soil salt-extractable organic carbon (SEOC)-a proxy for biotically-derived organic acid declined from the surface to 40-60 cm and then increase to its peak at 60-90 cm under CT, at 90-120 cm under NTNS and at 120-150 cm under NTSM (Fig. 1b). The NTSM increased the SEOC concentration at almost all soil layers compared with the CT and the NTNS (Fig. 1b), in which at the surface and 120-150 cm depth the contents of SEOC with NTSM were twice higher than CT. The relative contributions of SEOC to SOC (SEOC/SOC) in the NTSM were also always higher than in the CT and NTNS (Fig. 1c). Based on the estimated root depths and soil bulk density, total soil inorganic nitrogen available for the coming growing season in the NTSM and the NTNS was approximate 427.34 and 352.34 kg ha^−1^, respectively, while only 179.63 kg ha^−1^ in conventional tillage.

**Fig 1.**
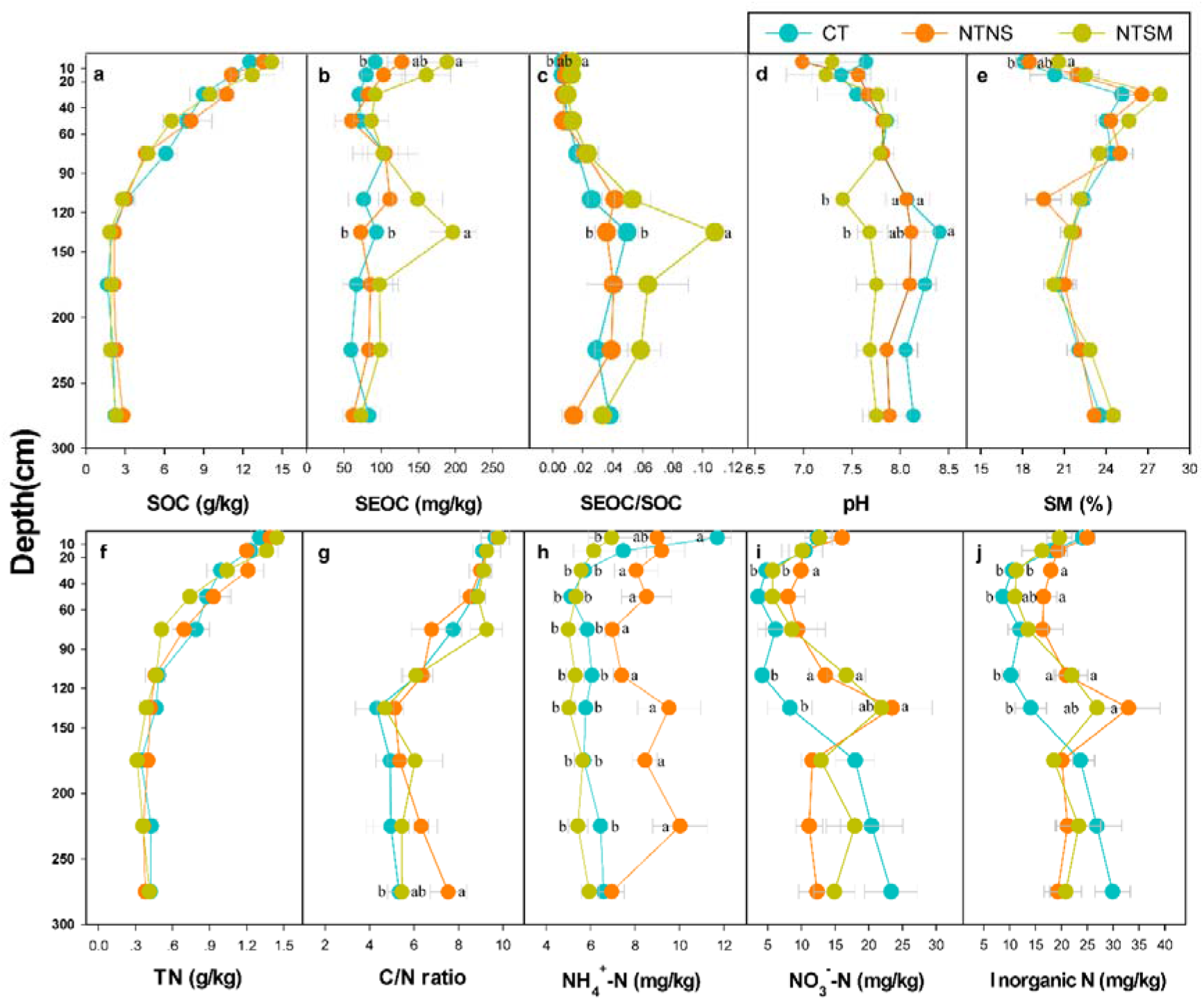
Soil properties (mean± SE, n = 3) along soil depth under different practices. **a**, SOC = soil organic carbon; **b**, SEOC = salt-extractable organic carbon; **c**, SEOC/SOC = ratio of SEOC to SOC; **d**, soil pH; **e**, SM = soil moisture; **f**, TN = total nitrogen content; **g**, C/N = ratio of SOC to TN; **h**, NH_4_ ^+^-N = ammonium nitrogen; **i**, NO_3_^−^-N = nitrate nitrogen; **j**, Inorganic N = NH_4_ ^+^-N + NO_3_ ^−^-N. Error bars indicate standard errors (n = 3). Different letters indicate significant differences (P < 0.05) among disturbing practices.

The mean annual corn yield (2013-2016) in the NTSM is 13416.8 kg/ha, which is much higher than the CT and NTNS (Fig. 2), particularly during the drought year of 2015, with only 409.6 mm of rainfall during the growing season (about 100 mm lower than the mean rainfall), while the corn yield in NTSM is 36.4% and 22.3% higher than the CT and NTNS, respectively (Fig. 2).

**Fig 2.**
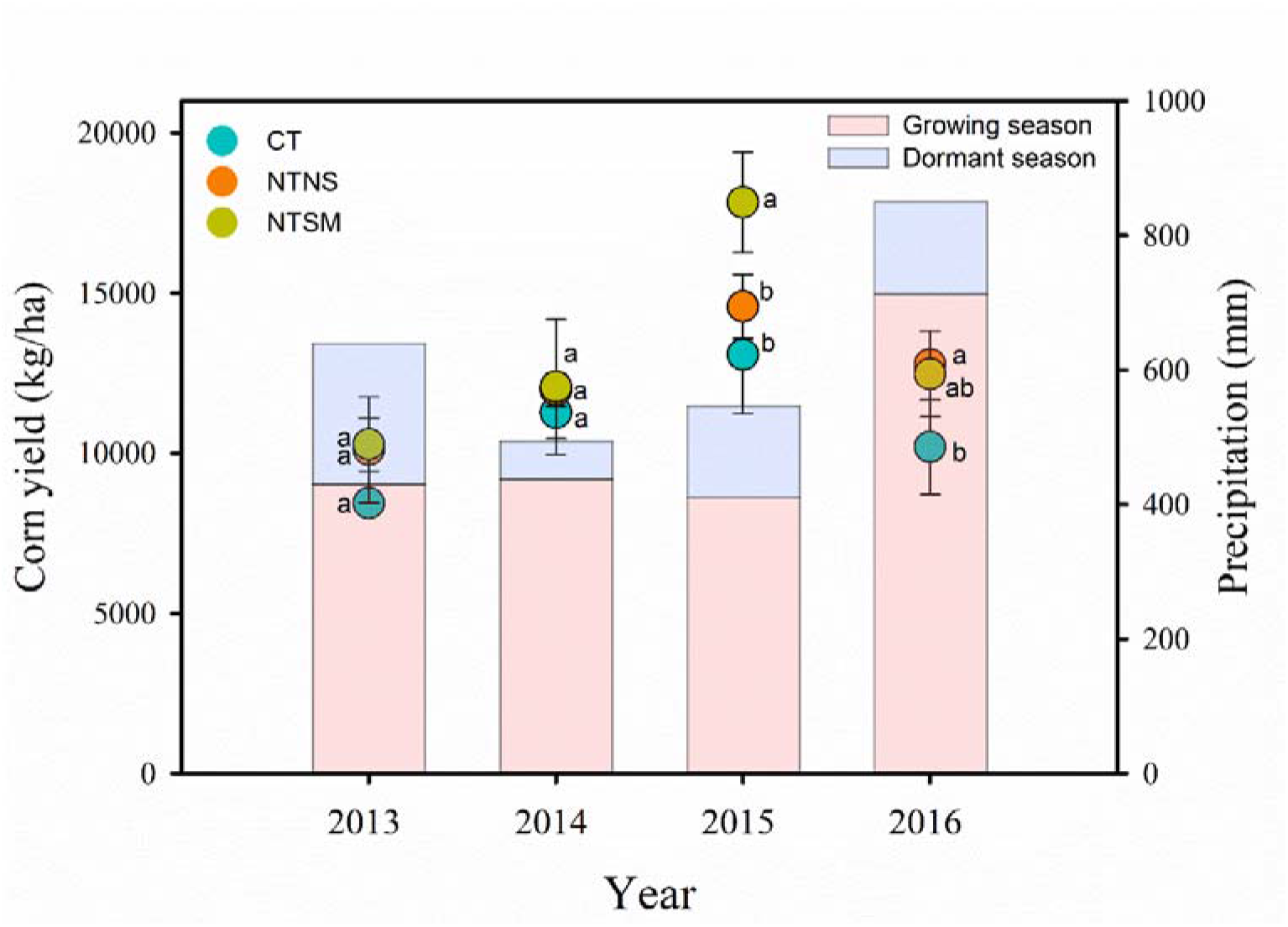
Corn yield (line+ symbol) and annual rainfall during growing and dormant seasons (bar) under different disturbance practices during 2013-2016. Error bars indicate standard errors (n = 3 or 4), different letters indicate significant differences at P < 0.05. Different corn seeds were used among years.

### Microbial diversity, composition, and structure

The microbial richness (Chao1), observed number of species (Observed-species) and diversity (Shannon-Index) first increased within 0-20 cm and decreased from 20 to 90 cm, then increased hereafter (Fig. 3). The low-disturbance practices significantly increased Chao1, Observed-species and Shannon-Index, particularly in 0-40 cm soil depths (Fig. 3). There were 54 microbial phyla across all soil samples. The dominant phyla (relative abundance > 1% across all soil samples) were Proteobacteria, Actinobacteria, Chloroflexi, Acidobacteria, Nitrospirae, Gemmatimonadetes, Planctomycetes, and these phyla accounted for 60-91% of the total microbial abundances in the whole soil profile (Fig. S2a). Bacteroidetes, Verrucomicrobia, Latescibacteria, Parcubacteria, Firmicutes, Microgenomates and Saccharibacteria were less dominant (relative abundance > 0.1% across all soil samples) but were still found across all soil samples (Fig. S2a). Although no difference in the composition of dominant phyla among treatments was found, there were more non-dominant phyla with higher relative abundance in low disturbance practices than conventional tillage practice (Fig. S2b).

**Fig 3.**
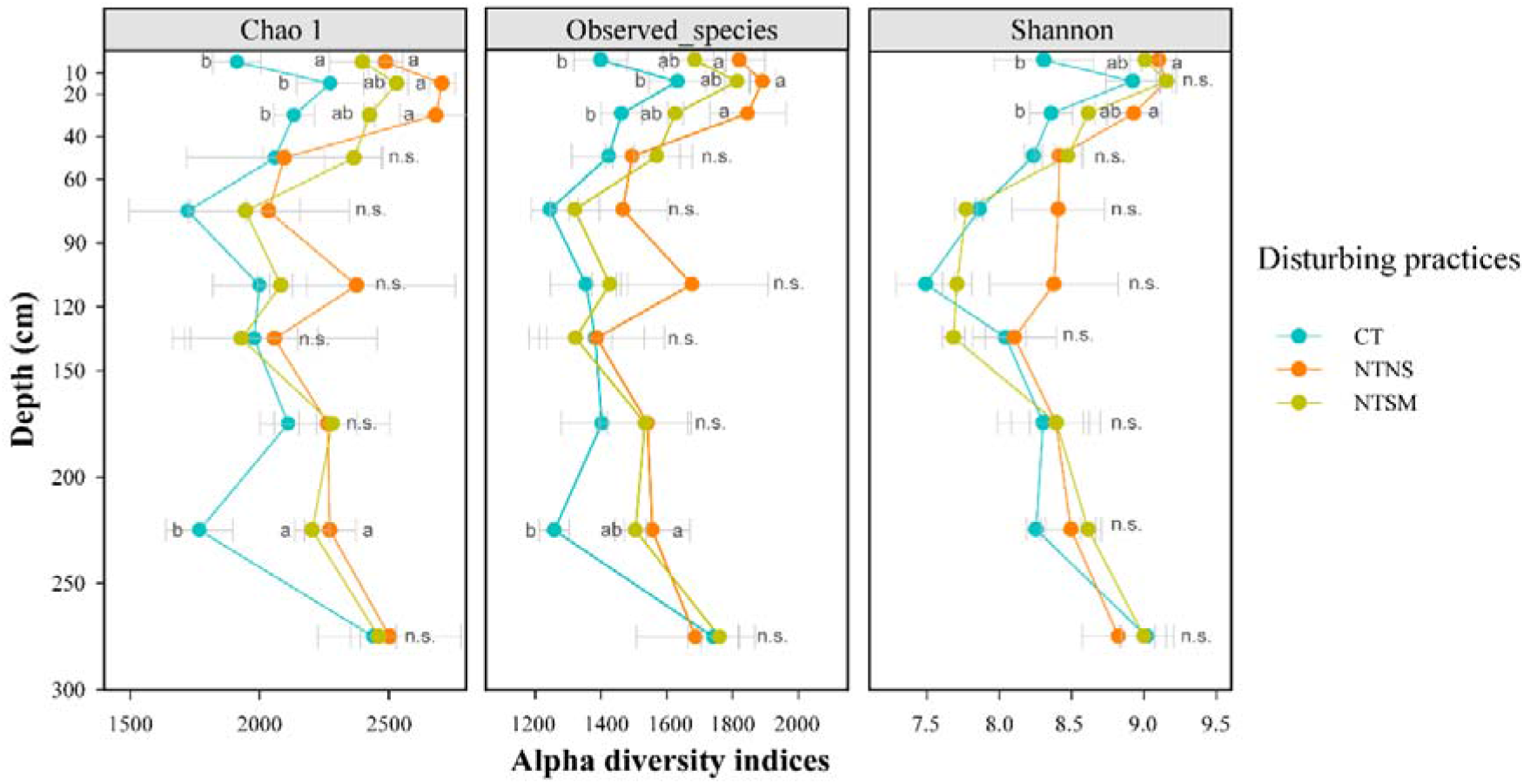
Microbial richness (Chao1) observed number of species (Observed_species) and diversity (Shannon_Index) in the CT, NTNS and NTSM plots. Error bars indicate standard errors (n = 3). Different letters indicate significant differences (P < 0.05) among disturbing practices.

Indicator analysis identified 16 and 51 clearly classified genera (relative abundances > 0.005%) in the NTNS and the NTSM plots, respectively, while no indicator genera were found in the conventional tillage plots (Fig. 4 and Table S2). The indicator genera in the NTNS plots belonged to Proteobacteria, Actinobacteria, Chloroflexi, Gemmatimonadetes and Planctomycetes, and most of them appeared in the surface soil (0-20 cm) with only 1 genus below 150 cm. Importantly, more extra indicator genera − including Bacteroidetes, Acidobacteria, Deferribacteres, Firmicutes, Verrucomicrobia, Chlorobi and Spirochaetae - existed in the NTSM plots, in which under 150 cm we observed 7 genera (Fig. 4 and Table S2).

**Fig 4.**
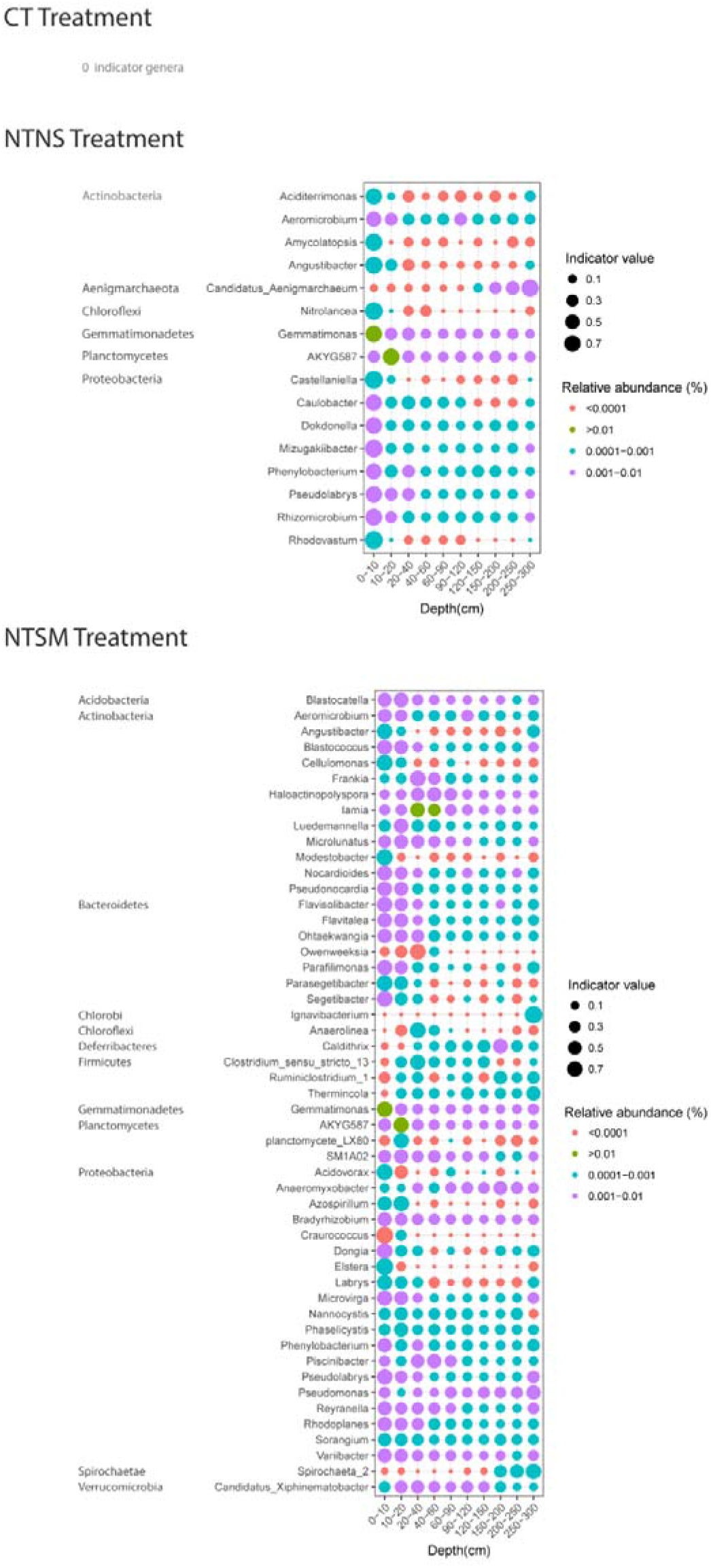
Indicator genera significantly (q < 0.1) associated with tillage practices. The size of each circle represents the indicator value of a specific genus in the different soil depths. The color indicates the relative abundance of each indicator genus. Taxonomic information, indicator values, P-values, and q-values of all indicator genera are given in Table S2. Zero indicator genera were identified in CT treatment.

Microbial community structures were visualized by Non-metric multidimensional scaling (MDS) and tested by PERMANOVA based on Bray–Curtis. The microbial communities among treatments in the root zones were marginally different (PERMANOVA p=0.08); however, below the root zone they differed distinctively (PERMANOVA p=0.02). The disturbance practices influenced the vertical distribution dissimilarity in microbial community structure (Fig. 5). Three clusters − 0-10 cm and 10-20 cm, 20-150 cm and 150-300 cm - were observed in the CT plots (PERMANOVA-F=9.57, p=0.0001) (Fig. 5). In the NTNS plots, 0-10 cm formed an independent cluster, while other soil depths showed some separation (e.g. 20-120 cm were separated from 150-300 cm soil depths by axis 1); however, Bray-Curtis distances between adjacent depths were too close to be separated (PERMANOVA-F=8.18, p=0.0001) (Fig. 5). The NTSM treatment clustered 0-10 cm and 10-20 cm together, 120-150 cm, 150-200 cm, 200-250 cm and 250-300 cm separately, and the other depths show some separations as well (PERMANOVA-F=11.32, p=0.0001) (Fig. 5).

**Fig 5.**
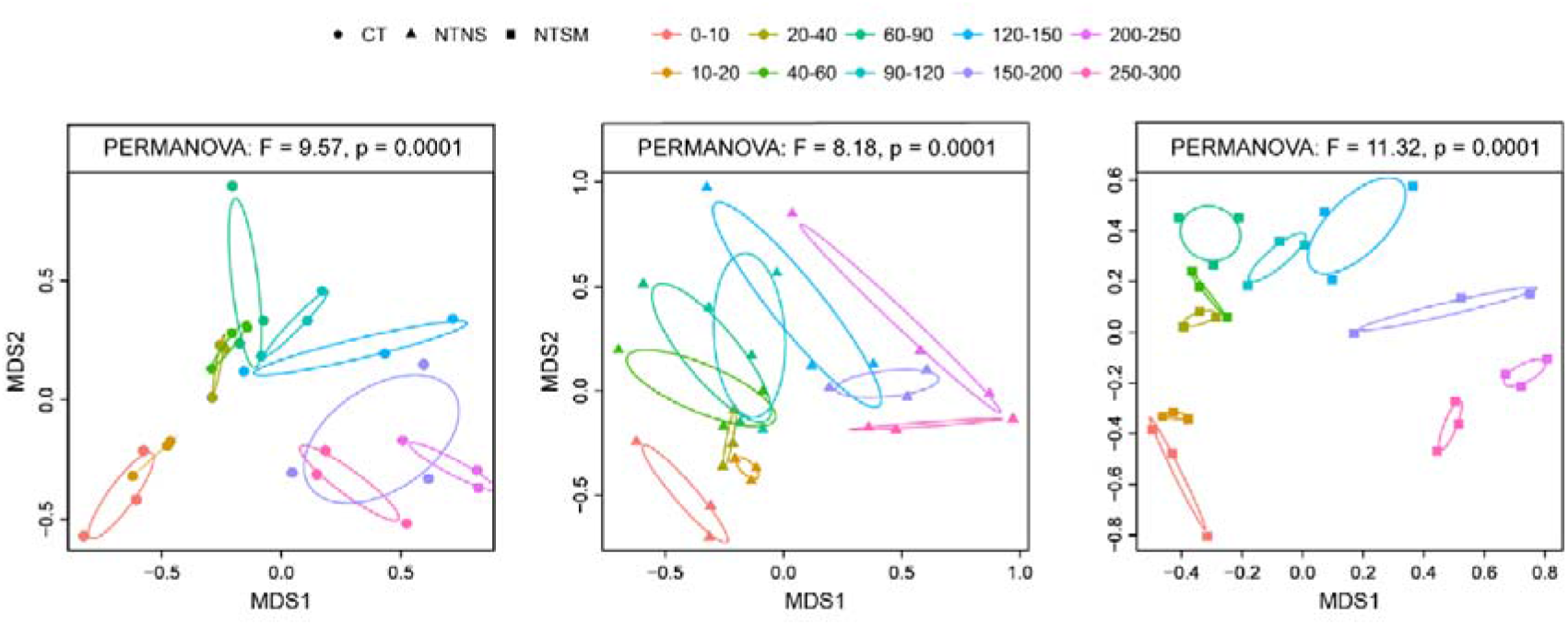
Non-metric multidimensional scaling (MDS) ordination of soil microbial community structures based on Bray-Curtis distances among soil depths at different agricultural disturbance practices. Permutational multivariate analysis of variance (PERMANOVA) revealed that the overall microbial community structures among soil depth were significantly different at each disturbance practice. Circles, triangles and squares represent CT, NTNS and NTSM, respectively.

### Predicted Ecological functions of microbial communities

According to the results of microbial diversity, composition and structure, the metabolic capabilities of microbial community in the whole 3-m soil profiles were predicted using Tax4Fun (Fig. S3). Results showed that low-disturbance practices significantly increased the abundance of predicted functions related to carbohydrate metabolism, nucleotide metabolism, glycan biosynthesis and metabolism, lipid metabolism and metabolism related to cofactors and vitamins (Fig. S3a). Moreover, the relative abundances of genes encoding for assimilatory nitrate reduction in low-disturbance practices were higher than that in conventional tillage practice (Fig. S4). The results suggested that in low disturbance practices, microbial community prefer to convert the nitrate/nitrite to ammonia. We then further assessed the impact of stover mulching on functional profiles (Fig. S3b). The extended error bar plot shows that the NTNS enriched the abundance of amino acid metabolism and lipid metabolism, while the NTSM enriched the functions associatedto energy metabolism, carbohydrate metabolism, biosynthesis of secondary metabolites, glycan biosynthesis and metabolism as well as metabolism of cofactors and vitamins (Fig. S3b).

### Relationships between microbial communities and soil properties

Forward selection in Redundancy analysis (RDA) revealed that soil depth (pseudo-F=48, p= 0.002), SOC (pseudo-F=11.5, p= 0.002), SM (pseudo-F=3.4, p= 0.012), soil pH (pseudo-F=2.3, p=0.018) and soil NH_4_^+^-N (pseudo-F=2.7, p= 0.026) significantly affected the vertical distribution of microbial communities (Fig. S5). Furthermore, the soil properties that regulated the distribution of soil microbes were different under different disturbance practices. Under the CT treatment, soil microbial community was mainly affected by soil NH_4_^+^-N (pseudo-F=4, p= 0.002) and soil NO_3_^−^-N (pseudo-F=2.3, p= 0.012) that mainly came from applied fertilizer (Fig. 6). The microbial community positively correlated to soil NH_4_^+^-N in the 0-20 cm soil, to soil NO_3_^−^-N negatively within 20-150 cm, while to soil NO_3_^−^-N positively after 150 cm (Fig. 6). Under the NTNS treatment, soil pH (pseudo-F=3.7, p=0.004) constrained the distribution of the microbial community, in which strong negative correlations occurred in 0-10 cm soil and a positive correlation in 90-150 cm (Fig. 6). Under the NTSM treatment, soil TN (pseudo-F=11, p=0.002), SM (pseudo-F=2.6, p=0.004) and C/N ratio (pseudo-F=1.8, p=0.016) significantly influenced the soil microbial community separation (Fig. 6). In general, the microbes positively correlated with the soil TN and C/N ratio in the surface soil layers (0-40 cm) and with SM in the middle layers (40-150 cm), while they were mainly influenced by depth in the deeper soil (150-300 cm) (Fig. 6).

**Fig 6.**
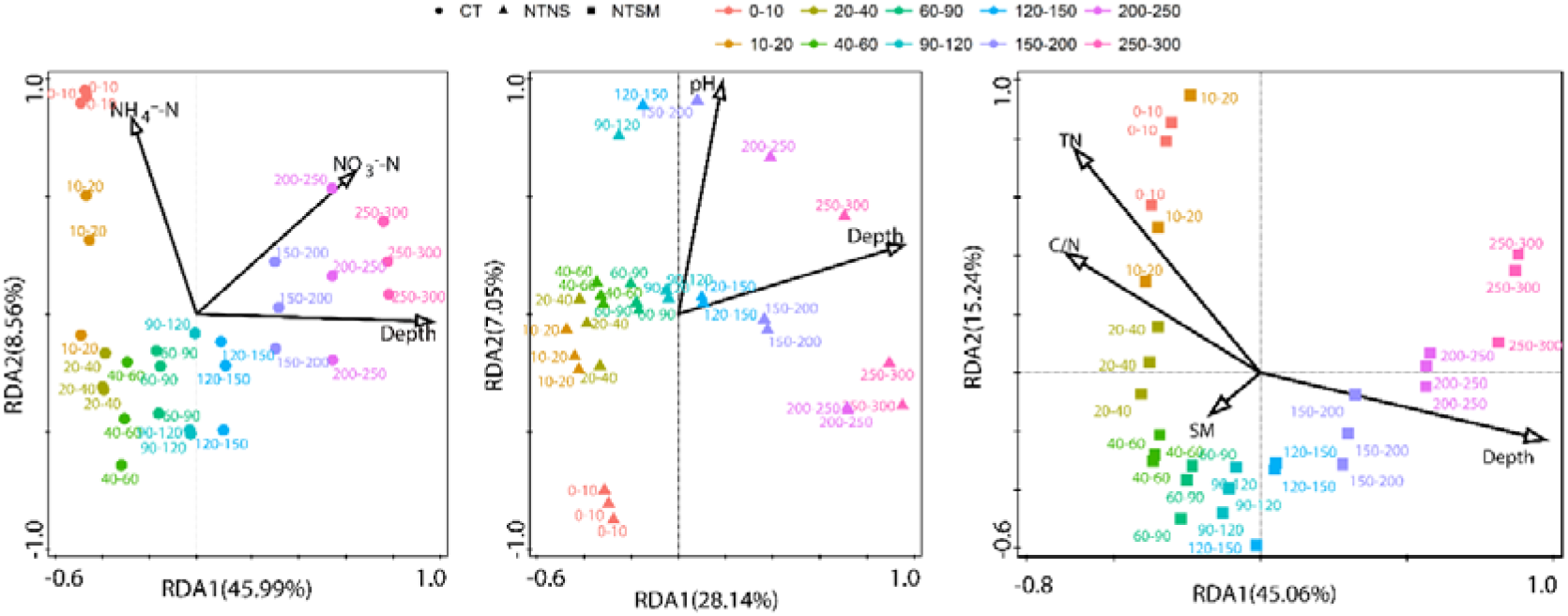
Redundancy analysis (RDA) of the soil microbial community originating from microbial phyla constrained by soil properties under different agricultural practices. Only soil variables that significantly explained variability in microbial community structure in the forward selection procedure were selected to the ordination (arrows). TN, total nitrogen content; C/N, a ratio of carbon to nitrogen content; NH_4_^+^-N, ammonium nitrogen; NO_3_^−^-N, nitrate nitrogen;SM, soil moisture. Circles, triangles and squares represent CT, NTNS and NTSM, respectively.

### Root depth among tillage practices

Currently the measurement of fine roots with diameter less than 0.2 mm are still technically difficult and their high turnover rates in-situ make the measurement even more complex(Pierret *et al*., 2016). Instead of detecting fine roots directly, we measured SEOC-a proxy for biotically-derived organic acid, which is a sensitive signal of root density and could be an indicator of root depth(Billings *et al*., 2018). We also did a literature review on whether no-tillage and straw mulching extend root depth in other crops. According to the change of SEOC with soil depth under different disturbing practices(Fig. 1b), we estimate that corn roots reached up to 60-90 cm, 90-120 cm and 120-150 cm under the CT, the NTNS and the NTSM, respectively, which is in line with reported corn root depths (∼150 cm) (Canadell *et al*., 1996; Kemper *et al*., 2011).According the literature review, we can’t find root-depth studies including both no-tillage and straw mulching. However, no-tillage as a conservative management increases root depth by 23 cm, 39 cm and 14 cm in corn, wheat, and sunflower, respectively (Fig. S6).

## Discussion

### No-tillage practices promote soil nutrient, water holding capacities and corn yield

The changes of SEOC with soil depth and the literature review of root depth among tillage indicated that no-tillage could promote root growth into deep soil, up to 150 cm in the NTSM. The root exudates with various organic acid and dead roots likely contributed to the lower soil pH in the NTSM, as shown by a significant negative relationship between SEOC and pH (r=–0.678, p<0.05). Despite function predication analyses have various uncertainty, the increased dissimilatory and assimilatory nitrate reduction genes suggested that low disturbance practices tend to convert nitrate to ammonium (Fig. S4), while lower ammonium in NTSM may be due to ammonium subsequently incorporated into biomass, as hints by the increase in genes related to biosynthesis metabolism and glycometabolism (Fig. S3). Moreover, maize stover or increased fine roots in deeper soil in NTSM provided labile carbon (Fig. 1b) to remove leaked nitrate through denitrification in deeper soil (below 1.5 m), where higher relative abundance of the denitrification bacteria (*Pseudomonas* and *Caldithrix*)(Koike and Hattori, 1975; Miroshnichenko *et al*., 2003) (Fig. 4 and Table S2) were detected. However, shallower roots in the CT treatment can’t provide enough labile carbon to remove extra soil NO_3_^−^-N in deep soil, thus causing nitrite accumulation and leaching into deeper soil layers. The amount of inorganic nitrogen accumulated in the root zones under NTSM (427.34 kg ha^−1^) likely could provide plenty of nitrogen for corn growth in the coming growing season (Fig. 2), based on the removed nitrogen in the grain (∼200 kg ha^−1^). Additionally, in line with many studies that show stover mulching reduces water evaporation and surface runoff and increase soil moisture in top soils(De Vita *et al*., 2007; Prosdocimi *et al*., 2016), we found that the topsoil moisture was significantly higher in the NTSM than in the CT plots (Fig. 1e) when the precipitation during the entire dormant period was only 38.5 mm. More importantly, the topsoil (0-20 cm) SOC stock in NTSM (18.5 Mg/ha) were significantly higher than in CT (15.8 Mg/ha) due to slightly higher bulk density in NTSM. Therefore, no-tillage with stover mulching not only increased SOC stock, the holding capacities for nutrients and water, thus reducing energy input to farm, but also tended to reduce the risk of nitrate leaching into groundwater. And more importantly, the healthy deep soil in turn raises corn production and promote the crop resistance to drought (Fig. 2). All these are critical to the development of sustainable agriculture and the associated ecosystems.

### No-tillage with stover mulching promotes microbial diversity, richness, and ecological function contributing to sustainable farming

Under the CT treatment, tillage heavily disturbed the topsoil and liberated occluded organic materials. Microbes tended to rapidly use available nutrients in the plowed layer (e.g. NH_4_^+^-N)(Ramirez-Villanueva *et al*., 2015), thereby causing the reduction of microbial metabolic diversity (Fig. S3a). Then, the resistance of the soil to stress or disturbance may also decrease(Kremen, 2005). In deeper soil layers, due to shallower roots, NO_3_^−^-N could quickly move downward and accumulate in deeper soil (Fig. 1i), which not only contaminated the underground water but also limited the activity of non-dominant microbes with important ecological functions, as no indicator genera were identified for each soil depth in CT treatment (Fig. 4 and Table S2). Because the microbial communities were closely associated with inorganic nitrogen, the microbes under CT were mainly influenced by added chemical fertilizer(Wood *et al*., 2015). Although the dominant microbial communities in CT were similar to those in the NTNS and NTSM, the loss of function resulted from the difference of non-dominant microbes, indicating that the soil under CT had degraded.

Under the NTNS treatment, soil pH was the major edaphic factor affecting the microbial community and the indicator genera (Fig. 6 and Table S3). The lower soil pH possibly was caused by deeper roots as shown by higher SEOC that is generally positively related to root density(Billings *et al*., 2018). Soil pH is often observed as a major factor determining the microbial composition and structure in natural ecosystems(Lauber *et al*., 2009; Zhalnina *et al*., 2015), as microbes often show a narrow tolerance to soil pH. In addition, soil pH regulates the availability of nutrient and mitigate ion toxicity(Lauber *et al*., 2009; Rousk *et al*., 2010; Zhalnina *et al*., 2015). Under NTNS, soil pH and depth explained 35% distribution of the microbial community (Fig. 6). We speculated that other edaphic factors (e.g. salinity and iron) directly or indirectly related to soil pH and SEOC also influenced the changes in the microbial community.

Under NTSM treatment, TN and C/N significantly correlated with soil microbial community due to the high C/N ratio of stover or roots (Fig. 6). Prior studies have reported that, following maize stover mulching, more organic N, amino acid N, and amino sugar N were observed in soil(Liu *et al*., 2016; Lu *et al*., 2018), combining with higher genes related to amino acid metabolism and carbohydrate metabolism from function predication analyses (Fig. S3), jointly hint that NTSM increased the retention time of nitrogen, hence meeting the nutrient requirement of corn growth and reducing nitrate loss to underground water. The increased available nitrogen, labile carbon and water in deep soil under NTSM can increase the resilience and resistance of maize to disturbances with higher grain production (Fig. 2). Zhang et al.(Zhang *et al*., 2013a) also observed litter-covered soil showed greater resistance to heating and copper addition due to the changes in soil properties and microbial community structure. Resistance to disturbance or stresses is the nature of a healthy soil and is essential for maintaining ecosystem functions, such as decomposing organic matter(Kibblewhite *et al*., 2008; Zhang *et al*., 2013a). Under the NTSM treatment, the microorganisms associated with the degradation of relatively stable carbon compounds, such as Planctomycetes and Verrucomicrobia (Table S4)(Herlemann *et al*., 2013; Erbilgin *et al*., 2014) as well as the indicator *Cellulomonas* and *Azospirillum* (Fig. 4 and Table S2) with the function of cellulose decomposition(Pathma *et al*.; Halsall and Goodchild, 1986) were increased. The predicted functional profiles related to energy metabolism (Carbon fixation pathways in prokaryotes), carbohydrate metabolism (TCA cycle, amino sugar, nucleotide sugar, galactose, fructose), biosynthesis of secondary metabolites (Carotenoid and Betalain) and glycan biosynthesis were increased, suggesting a higher metabolic activity and a change in substrate quality (Fig. S3). In addition, stover mulching also increased the ecological filter function of soil depth for selecting microbial communities as more indicator genera of each soil depths were identified under NTSM compared to NTNS and CT practices (Fig. 4 and Table S2). And these indicators residing at different soil depths might enhance the anti-disturbance ability of NTSM. For example, denitrification bacteria *Caldithrix* and *Pseudomonas*(Koike and Hattori, 1975; Miroshnichenko *et al*., 2003) were the indicator genera of 150-200 cm and 250-300 cm, respectively (Fig. 4 and Table S2), which might explain the low nitrate in the deep soil in NTSM. *Ignavibacteria* and *Spirochaeta*, the indicator genera of deep soil, have the ability to grow under the conditions of strictly anaerobic(Iino *et al*., 2010) and severely limited nutrients(Terracciano and Canale-Parola, 1984), respectively. Surface indicator genera belonging to Bacteroidetes might have the ability to degrade organic matter that is difficult to decompose(Thomas *et al*., 2011).

### Implications for climate change and food security

It was observed that about 179.63, 352.34 and 427.34 kg ha^−1^ inorganic N were kept in the root-zone soil in the CT, NTNS and NTSM, respectively. Generally, corn roots reach their maximum depth at the silking stage(Archontoulis and Licht, 2017), which is also the time when the heaviest rainfall occurs in northeastern China. We therefore expect that the available N kept in the root zone would be utilized by crops in the coming growing season before the storm leach the nitrogen down to ground water, which means that fertilizer N could be cut to meet crop growth in, at least, Northeast China and also prevent reactive N losses. Since the nitrogen use efficiency (NUE) of maize system under the conventional management is 51% in northeast China (NUE is defined as the efficiency of fertilizer N transferring to harvested crop N)(Zhang *et al*., 2019). Then, we conservatively calculate the required fertilizer N in the next year based on two assumptions: 1) the NUE of soil available N in root zone is equal to that NUE of applied fertilizer N, both of them are 50%; 2) the mineralized N during the coming growing season is neglected. Thus, N supply requirement = Fertilizer N×NUE + N in root zone ×NUE + Stover-N, where Stover-N for NTSM is 60 kg ha^−1^. We estimated the N requirement for each disturbance practice by multiplying grain yield by grain N concentration (1.4%)(Zhang *et al*., 2019) plus multiplying stover yield by stover N concentration (0.8%)(Izewska and Woloszyk, 2015). For CT, NTNS and NTSM, the mean annual corn yields were 10946.74, 12487.81 and 13416.81 kg ha^−1^, and the stover yields were 966.67, 10083.33 and 10833.33 kg ha^−1^, respectively. Thus, the N requirements were 230.6, 255.5 and 274.5 kg ha^−1^ for CT, NTNS and NTSM, respectively. Therefore, the theoretically conservative amounts of fertilizer N in the coming growing season are 281.6, 158.7 and 1.7 kg ha^−1^ for CT, NTNS and NTSM, respectively. No fertilizer-N is needed to apply without reducing corn yield in the NTSM plot. Compared to CT, the NTNS and NTSM could at least save respectively about 122.9 and 281.6 kg ha^−1^ N-fertilizer. For every kilogram of fertilizer-N produced and used on cropland, up to 87.9 MJ of energy is consumed(Kennedy, 2000) and 13.5 kg of CO_2_-equivalent (eq) (CO_2_-eq) is emitted(Zhang *et al*., 2013b). Hence, totally 24,752.6 MJ of energy consumption could be reduced and 3,801.6 kg CO_2_-eq emission could be cut per hectare cornland in Northeast China at least by using NTSM tillage practice. If this could be applied to all maize farmland in Northeast China (13,000,000 ha, Source: China Statistics Yearbook 2018), 0.3 EJ of energy could be saved and 49.4 Mt of CO_2_-eq could be reduced. Based on the average annual energy consumption for households of China in 2017 (15 EJ, China Statistics Yearbook 2018) and CO_2_ emissions (9,839 Mt, Global Carbon Atlas), the NTSM practice in corn farming of Northeast China has the potential to save 2% of household energy and to reduce 0.5% of CO_2_ emissions each year in the whole country.

Our results, particularly higher SEOC content, microbial diversity, and indicator genera in the NTSM deep soil compared with CT, clearly showed that low-disturbance practices can affect agricultural resource in deeper soil over time. This indicates that crops, like corn in this study, under appropriate management can ultimately explore nutrients and water from deeper soil, thus not only increasing the volume of soil (almost double in this study) exploited without reclaiming more natural land areas but also reducing nutrient loss into ground water. Meanwhile, the input of labile carbon including SEOC into deeper soil in the NTSM provides essential energy and nutrients to microbes and gradually shape highly diversified and functional microbial communities in the deep soil over time, and hence improve the self-sustaining ability of farmland in the face of climate change. The improvements of microbial communities in the deep soil (1-3 m) at the end of dormant season in our study provide evidence for the first time that a nature-based management in farmland is conducive to deep-soil health for sustainable farm in a long run. Although many scientists have realized the importance of deep rooting for sustainable intensification of crop production(Pierret *et al*., 2016; Thorup-Kristensen *et al*., 2020), the deep-root studies are still rare due to technological bottleneck. According the literature review, no-tillage extend root depth in other crops, like corn, wheat and sunflower. While some studies also show no effects or even reduce root depth(Dwyer *et al*., 1996; Izumi *et al*., 2004), the possible reason is the legacy effects of tillage, for example, long-term no-tillage leads to soil compaction, and the soil system might be still in its transition stage. Coupled with our results, low-disturbance practices as a nature-based agricultural management likely can develop a deeper root system to explore more resource in deep soil to sustain food production, while without threatening environmental security and reclaiming more lands.

## Materials and Methods

### Site description and soil sampling

The field experiment was established in 2007 at the Lishu Conservation Tillage Research and Development Station of the Chinese Academy of Sciences in Jilin province, Northeast China (43.19° N, 124.14° E). The region has a humid continental climate with a mean annual temperature of 6.9 °C and the mean annual precipitation of 614 mm. The soils are classified in the Mollisol order (Black Soil in Chinese Soil Classification) with a clay loam texture(IUSS Working Group, 2006). The site has been continuously planted with maize since 2007. We set up an experiment by a randomized complete block design with four replicates and five treatments. Each plot area was 261m^2^ (8.7×30m). The five treatments included conventional tillage (moldboard plowing to a depth around 30 cm and removed the stover), no-tillage (no soil disturbance and direct seeding), and no-tillage with three-level stover mulching (33%, 67% and 100% newly produced maize stover were evenly spread over the soil surface each fall) (Fig. S1). For each treatment, slow-release fertilizer was applied at one time when sowing, which was equal to 240 kg/ha N; 47 kg/ha P; 90 kg/ha K. The rainfall data were obtained from local meteorological administration. The grain yield was estimated by manually harvesting 20 m^2^ area, randomly taken from each plot.

In this experiment, in order to reduce the damage to the plots and reduce costs, 3 plots were randomly taken from each treatment including conventional tillage (CT), no-tillage without stover mulching (NTNS), no-tillage with 100% stover coverage (NTSM) as three comparative practices. In April 2017, triplicate soil cores (0-300 cm) were collected from each plot at the end of dormant season. After removing surface stover, we took soil cores by a stainless-steel hand auger and sliced each into ten layers: 0-10 cm, 10-20 cm, 20-40 cm, 40-60 cm, 60-90 cm, 90-120 cm, 120-150 cm, 150-200 cm, 200-250 cm, 250-300 cm. In total, 90 soil samples were collected and transported to the laboratory within 3 hours, then passed through a 2-mm sieve. All visible roots, crop residues and stones were removed. Each soil sample was divided into three subsamples: one subsample for DNA extraction and soil salt-extractable organic carbon (SEOC) measurement that was immediately placed into a polyethylene plastic bag and stored at −80 °C, one for chemical measurements including ammonium nitrogen (NH_4_^+^-N) and nitrate nitrogen (NO_3_ ^−^-N) (within one day), and the remaining one was air dried for other soil physicochemical properties.

### Soil properties

Soil total nitrogen (TN) content was measured by an Element analyzer Vario EL III (Elementar Analysensysteme GmbH, Hanau, Germany). Soil organic carbon (SOC) was converted from soil organic matter that was measured by potassium dichromate oxidation(Nelson and Sommers, 1982). Soil pH was measured in deionized free-CO_2_ water (1:2.5 w/v). Gravimetric soil moisture was determined by oven-drying fresh soil to a constant weight at 105 °C. Soil NH_4_^+^-N and NO_3_^−^-N were extracted from fresh soil by 2 M KCl and measured by a continuous flow analytical system (AA3, SEAI, Germany). To reflect soil soluble, exchangeable, mineral-bound OC, SEOC was extracted from the frozen soil samples with 0.5 M K_2_SO_4_ (1:5 w/v)(Jones and Willett, 2006; Toosi *et al*., 2012).

### DNA extraction, PCR amplification and pyrosequencing

Soil DNA was extracted from the frozen soil samples (0.5 g wet weight) by using MoBio PowerSoil DNA isolation kit (MoBio Laboratories, Carlsbad, CA, USA) following the instructions of the manufacturer. The quality of DNA was determined by 1% agarose gel electrophoresis. The V3–V4 region of the bacterial 16S rRNA gene was amplified by PCR using the primers 338F and 806R with barcode for Illumina MiSeq sequencing. PCR was performed in a total volume of 50 μl containing 30 ng DNA as a template, 20 mol of each primer, 10mM dNTPs, 5μl Pyrobest buffer (10×) and 0.3 U of Pyrobest polymerase (Takara Code: DR005A). Each sample was amplified for three replicates. The PCR products from the same sample were pooled, checked by 2% agarose gel electrophoresis and were then purified using AxyPrepDNA agarose purification kit (AXYGEN). Finally, purified PCR products were sequenced on an Illumina MiSeq platform PE300 sequencer (Illumina, USA).

The raw sequence data were further analyzed by the following protocol. Low-quality sequences with an average quality score of less than 20 were filtered by employing Trimmomatic(Bolger *et al*., 2014). The FLASH software was used to merge overlapping ends and treat them as single-end reads(Derakhshani *et al*., 2016). The non-amplified region sequences, chimeras and shorter tags were also removed using Usearch and Mothur(Mysara *et al*., 2016). The resulting high-quality sequences were clustered into Operational Taxonomic Units (OTUs) at 97% sequence similarity using Usearch (Version 8.1.1861 http://www.drive5.com/usearch/). OTUs were then classified against the Silva (Release119 http://www.arb-silva.de) database and the taxonomic information of each OTU representative sequence was annotated using the RDP Classifier(Wang *et al*., 2007). A total of 3,255,693 high-quality reads were obtained from all soil samples, which were clustered into 9,573 unique OTUs at a 97% sequence similarity. The Good’s coverage of all the samples ranged from 0.93 to 0.98, which indicates an adequate level of sequencing to identify the majority of diversity in the samples.

### Root depth data

Plant roots, fine roots in particular, release large amounts of labile organic carbon(Dakora and Phillips, 2002) that are essential for healthy microorganisms in soil. Generally, most studies considered that fine roots are roots < 2mm in diameter, while roots <0.2 mm in diameter can contribute to >50 % of the overall root length and play a major role in releasing root exudates and absorbing nutrients and waters(Pierret *et al*., 2016). Currently the measurement of fine roots with diameter less than 0.2 mm are still technically difficult and their high turnover rates *in-situ* make the measurement even more complex(Pierret *et al*., 2016). To minimize the root-turnover effects to the most degree, we collected soil samples at the end of dormant season, which can likely mirror the long-term legacy effects of our practices. Also instead of detecting fine roots directly, we measured soil salt-extractable organic carbon (SEOC)-a proxy for biotically-derived organic acid, which is a sensitive signal of root density and could be an indicator of root depth(Billings *et al*., 2018).

We also did a literature review on whether no-tillage and straw mulching extend root depth in other crops. We collected root depth of main crops data from literature published in 1996-2019 using the Web of Science resource and only the data with more than two tillage managements are retained. Finally, a total of 87 observations were selected (Table S5).

### Statistical analyses

Soil properties were analyzed and plotted using Sigmaplot 12.5 software. Alpha diversity indices were calculated in Qiime (version v.1.8) and used to reflect the diversity and richness of the microbial community in different samples. The relative abundances of individual phyla in different samples were computed by R packages. The indicator analysis based on genera-specific to each soil depth was conducted using indicspecies package of R with 9999 permutations, and the P-values were corrected for multiple testing using qvalue package of R(Cáceres and Legendre, 2009). Functional profiles of the microbial community were predicted by Tax4fun (an open-source package in R)(Aßhauer *et al*., 2015) and further statistical analysis was conducted by STAMP using Welch’s t-test (Parks *et al*., 2014). Non-metric multidimensional scaling (MDS) was performed by “vegan” package of R to describe differences in microbial community structure among samples. Permutational multivariate analysis of variance (PERMANOVA) was employed on Bray-Curtis distances to test the differences in soil microbial communities among various sample groups. The redundancy analysis (RDA, Canoco 5 software) were conducted to identify the correlations between microbial community composition and environmental variables. ANOVA were conducted by SPSS Version 22. Percentage data were transformed using arcsine square root function before ANOVA test. All statistical tests were significant at p ≤ 0.05.

## Sequence availability

All sequencing data that support the findings of this study have been deposited in the National Center for Biotechnology Information (https://www.ncbi.nlm.nih.gov/), in the Sequence Read Archive (SRA) database (BioProject number: PRJNA488172).

## Acknowledgements

We would like to thank Dr. William H. Schlesinger at the Cary Institute of Ecosystem Studies for his comments and Dr. Randy Neighbarger at Duke University for language editing. This work was supported by the “National Key R&D Program” (No. 2016YFD0800103, No. 2016YFD0200307) and the National Natural Science Foundation of China (grant number, 41671297). We would like to thank Pengshuai Shao, Xuesong Ma and many individuals for assistance with sample collection, processing and analysis.

## Author Contributions

H.X., XZ and C.L. designed the experiment, F.D. did field and lab measurements, F.D., H.W. and C.L analyzed data and wrote the manuscript, and all the authors discussed results and commented on the manuscript.

## Supplementary information

Supplemental information may be found online in the Supplemental Information section at the end of the article.

